# Sexual dimorphism in the social behaviour of *Cntnap2* KO mice correlates with disrupted synaptic connectivity and increased microglial activity in the anterior cingulate cortex of males

**DOI:** 10.1101/2023.06.20.545580

**Authors:** Matt S. Dawson, Kevin Gordon-Fleet, Lingxin Yan, Vera Tardos, Huanying He, Kwong Mui, Smriti Nawani, Zeinab Asgarian, Marco Catani, Cathy Fernandes, Uwe Drescher

## Abstract

A biological understanding of the apparent sex bias in autism is lacking. We have identified *Cntnap2* KO mice as a model system to help better understand this dimorphism. Using this model, we observed social deficits in juvenile male KO mice only. These male-specific social deficits correlated with reduced spine densities of Layer 2/3 and Layer 5 pyramidal neurons in the Anterior Cingulate Cortex, a forebrain region prominently associated with the control of social behaviour. Furthermore, in male KO mice, microglia showed an increased activated morphology and phagocytosis of synaptic structures compared to WT mice, whereas no differences were seen in female KO and WT mice. Our data suggest that sexually dimorphic microglial activity may be involved in the aetiology of ASD, disrupting the development of neural circuits that control social behaviour by overpruning synapses at a developmentally critical period.

## Introduction

Autism spectrum disorder (ASD) is a neurodevelopmental disorder encompassing impairments in social reciprocity, communication, and restricted, repetitive patterns of behaviour (Battle 2013, Willsey et al 2022). ASD is highly heritable and persists across the lifespan, with estimates suggesting that it affects roughly 1:57 of the population in the UK (Roman-Urrestarazu et al 2021).

Historically, ASD has been widely regarded as a disorder mainly affecting the male sex (Loomes et al 2017). An in-depth meta-analysis estimates the male-to-female ratio of ASD to be approximately 4:1, although this imbalance tends to be even higher (about 9:1) for individuals with normal or high IQ (Loomes et al 2017, Roman-Urrestarazu et al 2021, Werling & Geschwind 2013, Willsey et al 2022). However, there may also be a diagnostic gender bias, meaning that girls who meet criteria for ASD are at disproportionate risk of not receiving a clinical diagnosis (Loomes et al 2017). Nonetheless, males do appear to be at greater risk of developing ASD overall with several prominent theories to explain this apparent sex bias proposed. These include an extreme male brain (Baron-Cohen 2002), the existence of a female protective effect (Jacquemont et al 2014), and a female autism phenotype dissimilar to a male autism phenotype (Kopp & Gillberg 1992). Overall, though, the biology behind this sexual dimorphism in ASD remains poorly elucidated (Loomes et al 2017).

Hundreds of genes have been classified as ASD risk genes (https://www.sfari.org/resource/sfari-gene/). Predominantly, these genes have roles in the regulation of gene expression or neuronal communication, implicating both developmental and functional changes in ASD (Satterstrom et al 2020). Most of the genes affecting neuronal communication are involved in the control of synapse development and function, leading to the description of some forms of ASD as a ‘synaptopathy’ (Ebrahimi-Fakhari & Sahin 2015).

A cell type known to play a substantial role in the pruning of synapses in the developing brain is microglia (Faust et al 2021, Paolicelli et al 2011, Schafer et al 2012, Stevens et al 2007), the resident immune cells of the brain (Thion et al 2018). There is strong evidence that dysfunctional microglia– neuron interactions may play a role in the pathogenesis of ASD (Li & Barres 2018, Neniskyte & Gross 2017), indicating that a perturbed cross-talk between microglia and neurons could lead to an erroneous synapse refinement, disturbing the development of neural circuits, particularly those controlling social behaviour. Interestingly, microglia are sexually dimorphic from early postnatal development onwards as reflected in their divergent transcriptomes (Guneykaya et al 2018, Hammond et al 2019, Schwarz et al 2012) and are prominently involved in the development of neural circuits controlling sex-specific behaviour (McCarthy et al 2017).

One particular ASD risk gene is *CNTNAP2,* which encodes the CASPR2 protein, a member of the neurexin family and a synaptic cell adhesion molecule. *CNTNAP2* is one of the largest genes in the human genome and gene-linkage studies have identified *CNTNAP2* mutations in autistic individuals and families. Interestingly, in some individuals both *CNTNAP2* alleles are mutated, and are predicted to lead to a complete loss of the functional protein (Rodenas-Cuadrado et al 2016, Strauss et al 2006, Watson et al 2014, Zweier et al 2009). *CNTNAP2* is highly expressed in frontal lobe circuits in the developing human brain, and using functional neuroimaging, a relationship between frontal lobar connectivity and common genetic variants in *CNTNAP2* has been demonstrated (Scott-Van Zeeland et al 2010).

The upregulation of *Cntnap2* RNA expression coincides with the onset of synaptogenesis and circuit formation in the cortex (Gordon et al 2016, Kroon et al 2019, Sacai et al 2020). CASPR2 is expressed at both excitatory and inhibitory synapses but not by microglia (Betancur et al 2009, Gerrits et al 2020), and has been shown to stabilize already formed synapses, while having little synaptogenic activity itself (Anderson et al 2012, Gdalyahu et al 2015, Varea et al 2015). Mice mutant for *Cntnap2* recapitulate core behavioural deficits typical of ASD, including impairments in social behaviour and communication, as well as repetitive behaviours (Penagarikano & Geschwind 2012). Lack of CASPR2 leads to perturbed dendritic spine density, disrupted synapse function, an imbalance in excitatory-inhibitory (E/I) signalling and impaired neural oscillations (Anderson et al 2012, Lazaro et al 2019, Liska et al 2018, Scott et al 2019, Varea et al 2015, Zerbi et al 2018).

In addition, a combined rs-fMRI/c-fos expression study showed circuit-specific connectivity defects in *Cntnap2* mutant mice, including deficits in the Anterior Cingulate Cortex (ACC) (Choe et al 2022). The ACC represents a major hub of the limbic networks which has been regularly associated with the aetiology of ASD in humans (Apps et al 2016, Catani et al 2016). In animal models, a conditional knockout (KO) of another ASD candidate gene, *Shank3,* in the ACC of mice resulted in functional impairments of excitatory neurons and defects in social interaction. Re-expression of *Shank3* in the ACC in mice with a full *Shank3 KO* restored social interaction (Guo et al 2019, Monteiro & Feng 2017).

Previous analyses of the role of *CNTNAP2* have predominantly been carried out in male KO, or in a mix of male and female KO mice, and not stratified by sex (Anderson et al 2012, Lazaro et al 2019, Liska et al 2018, Scott et al 2019, Varea et al 2015, Zerbi et al 2018). However, there is a small but growing body of evidence suggesting that the effects of *Cntnap2* are partly sex specific. To our knowledge, there are two other studies utilising this mouse model that report females as more likely to be neurotypical even if they carry the mutations. Townsend et al. (2017) report that a disruption of CASPR2 expression in males, but not females, results in a decreased visually evoked activity in dorsal stream-associated higher visual areas. This decrease was not observed in adult females. Schaafsma et al. (2017) illustrate the sex specificity of *Cntnap2* by showing that an interaction between the *CNTNAP2* mutation and maternal immune activation (MIA) increase the expression of corticotropin-releasing hormone receptor 1 (Crhr1), which leads to deficits in social recognition, but only in male mice. In addition, there are reports in humans of how mutations in the *CNTNAP2* gene lead to sex-specific structural alterations on MRI (Udden et al 2017), while a genetic association study of *CNTNAP2* variants and developmental dyslexia found that two polymorphisms (rs3779031, rs987456) were associated with an increased risk of dyslexia in females but not males (Gu et al 2018).

We set out to further investigate this possible sexual dimorphism of *Cntnap2* KO mice. While male *Cntnap2* KO mice showed strong social deficits in behavioural assays, female *Cntnap2* KO mice did not. Our analyses further revealed that in the ACC of male, but not of female, *Cntnap2* KO mice there was a transient reduction in spine density of pyramidal excitatory neurons, and an increased engulfment and phagocytosis of pyramidal presynapses by microglia in male KO mice compared to male WTs, with no differences observed for female WT versus KO mice.

## RESULTS

### Juvenile male – but not female – *Cntnap2* KO mice show deficits in social behaviour assays

We analysed juvenile (5-weeks old) mice (Fig. 1) whose prefrontal cortex development corresponds roughly to that of young adolescent humans (Chini & Hanganu-Opatz 2021). Male and female *Cntnap2* KO mice were investigated separately in social interaction assays (Grayton et al 2013, McFarlane et al 2008), allowing fine-grained measurements of interactions between pairs of juvenile mice. We assessed the number of the following parameters: social sniffing (sniffing above the middle of the body including the head, snout and upper body region of the conspecific), anogenital sniffing (sniffing below the middle of the body of the conspecific, primarily around the anogenital region of the conspecific), following a conspecific (mouse moves near the other mouse, follows its movements without making direct contact with the mouse), and these three behaviours combined for a measure of total social investigation/interaction (Silverman et al 2010).

**Fig. 1.**
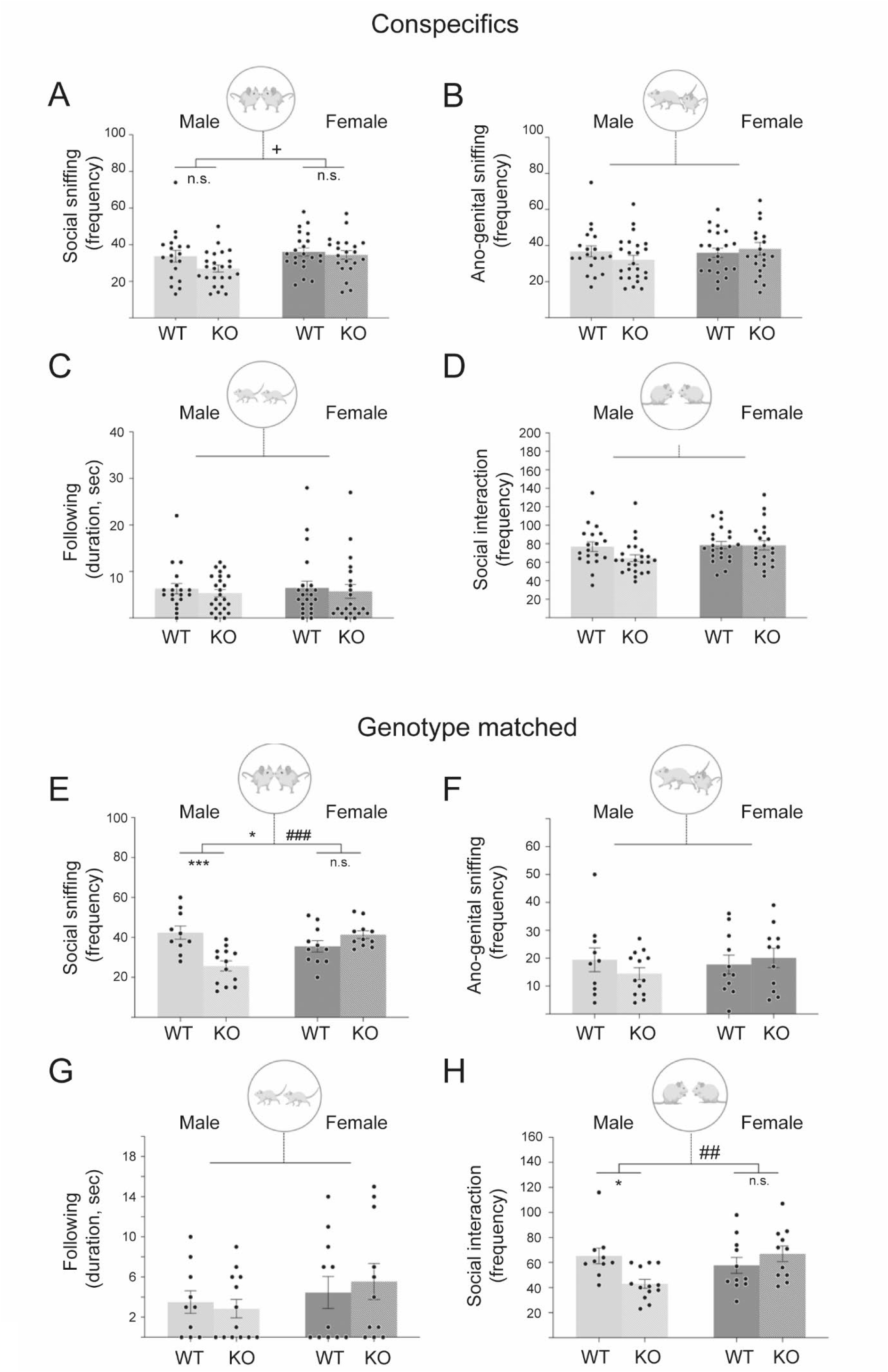
Behavioural deficits of juvenile male *Cntnap2* KO mice. Juvenile mice (5 weeks) were analysed in a social interaction assay, in which a test mouse (*Cntnap2* KO or WT mice of either sex) was placed in a novel cage, and a sex-matched juvenile conspecific C57BL/6J mouse **(A-D)** or an age-, sex-, and genotype-matched conspecific **(E-H)** was added. The frequency of their interaction was scored for 10 minutes for social sniffing **(A, E)**, ano-genital sniffing **(B, F)**, or following **(C, G)**. The total interaction frequencies are given in **(D,H)**. Single data points represent individual mice. Statistical analysis was performed using 2-way ANOVA. Symbols above the bars represent an overall effect of genotype (*), sex (+) or interaction (#), while symbols below the chart represent Tukey’s *post-hoc* significance between WT and *Cntnap2* KO (*). In each case 1, 2, or 3 symbols represents p < 0.05, p < 0.01, or p < 0.001. Juvenile male KO mice show significantly reduced frequency of social sniffing (E) and total social interaction (H), compared to WT mice. Data presented as means, error bars represent S.E.M. For A-D: n=20-26/group, for E-H: n=10-13/group. Full two-way ANOVA results can be seen in Table S1. Mouse images from Biorender.

We first assessed social investigation of test mice towards an unfamiliar, sex-matched juvenile conspecific C57BL/6J mice (Fig. 1A-1D) and then we measured social interaction between an unfamiliar, age-sex- and genotyped-matched pairs of test mice (Fig. 1E-1H).

For the social investigation of unfamiliar conspecifics (Fig. 1A-1D), two-way ANOVA revealed a significant effect of sex on the frequency of social sniffing (F1,82 = 4.2, p < 0.05), with female mice showing increased frequency of social sniffing compared to males but no effect of genotype There were no other significant genotype or sex differences for any of the other behavioural measures in the juvenile conspecific social investigation test.

When pairs of unfamiliar test mice matched by age, sex and genotype were tested in the social interaction assay, (Fig. 1E-1H), two-way ANOVA revealed a significant effect of genotype and interaction between genotype and sex on the frequency of social sniffing (F1,41 = genotype: 4.16, p < 0.05 and interaction: 17.79, p < 0.001). *Post-hoc* Tukey’s correction revealed a significant reduction in the frequency of social sniffing between male WT and male KO (p > 0.001), but not between female WT and female KO. The effect of genotype on social interaction was further seen in total social interaction frequency, with a significant genotype by sex interaction (F1,41 = 8.04, p = 0.007), post-hoc analysis revealed a significant reduction in the frequency of social interaction between male WT and male KO (p = 0.03), but not between female WT and female KO. There were no significant genotype, sex or interactions for any other behavioural measures taken in the social interaction tests.

All outcomes from behavioural statistical analyses and significant *post-hoc* corrections are documented in the supplementary index (Table S1).

Overall, juvenile male *Cntnap2* KO mice displayed reduced social behaviour when engaging in social interaction with an age-, sex- and genotype-matched unfamiliar mouse but not when investigating unfamiliar sex matched conspecific (C57BL/6J) mice. No behavioural deficits were seen between female WT and KOs, indicating that it is just male *Cntnap2* KO mice that display specific behavioural deficits.

There are a number of confounding factors that could affect mice behaviour so we also investigated these too. As social behaviour in rodents relies heavily on olfaction (Zou et al 2015), we investigated possible deficits in olfaction using the olfactory habituation/dishabituation test (Suppl. Fig. 1). We found normal habituation/dishabituation profiles in all groups (juvenile WT and KO mice of both sexes) – this is characterized by a decrease in sniffing latencies between the first and last exposure to a particular odour, followed by a re-instatement of sniffing when a new odour was presented (Suppl. Fig. 1). Our findings are in line with other investigations concluding that *Cntnap2* KO mice can discriminate between individual odours (Levy et al 2019).

We also used the open field assay to investigate other confounding factors including anxiety, and locomotor (hyper-) activity. Anxiety is an associated symptom of ASD, occurring in a subset of patients, and poses challenges regarding a discrimination between social anxiety and ASD proper (Silverman et al 2010). Our experiments showed similar anxiety levels for juvenile WT and KO mice of both sexes. In addition, we found similar locomotor activity and exploration in all groups (Suppl. Fig. 2).

### Transient decrease in apical dendritic spine density of pyramidal neurons in the ACC of male – but not female – *Cntnap2* KO mice

Social behaviour requires the processing of experiences from a diverse number of sources, including sensory and reward information as well as cognition and emotion (Gunaydin et al 2014). The ACC is anatomically and functionally connected to a broad set of regions engaged in social information processing and represents a crucial information hub of the ‘social network’ (Apps et al 2016).

Our aim was to investigate pyramidal projection neurons of Layer 2/3 (L2/3) and Layer 5 (L5) in the ACC, which appear to play an important role in controlling social behaviour by receiving and connecting to other intra-cortical and sub-cortical areas (Walum & Young 2018).

To determine whether loss of CASPR2 protein affects synapse densities during development in the ACC, we crossed *Cntnap2* KO with mice expressing the *grp:cre* transgene (Gerfen et al 2013) and a stop-floxed *tdTomato* expression cassette (*TOM^+^*) (jax.org). In *grp:cre*, *tom^+^* mice, tdTomato is expressed in a sub-population of pyramidal cells of L2/3 and L5 of the medial prefrontal cortex including the ACC (Fig. 2). This allowed analysis of individual fluorescently-labelled apical dendrites and spines projecting from these neurons in the outermost Layer 1 (L1). Layer 1 receives multiple inputs and also contains inhibitory interneurons, allowing it to orchestrate bottom-up and top-down signalling via L2/3 and L5 pyramidal neurons (Genescu & Garel 2021, Ibrahim et al 2020). Interestingly, many ASD candidate genes are expressed in this layer (Mayer et al 2018), suggesting that maladaptive integration of circuit development in this region raises susceptibility to the development of neuropsychiatric disorders.

**Figure 2.**
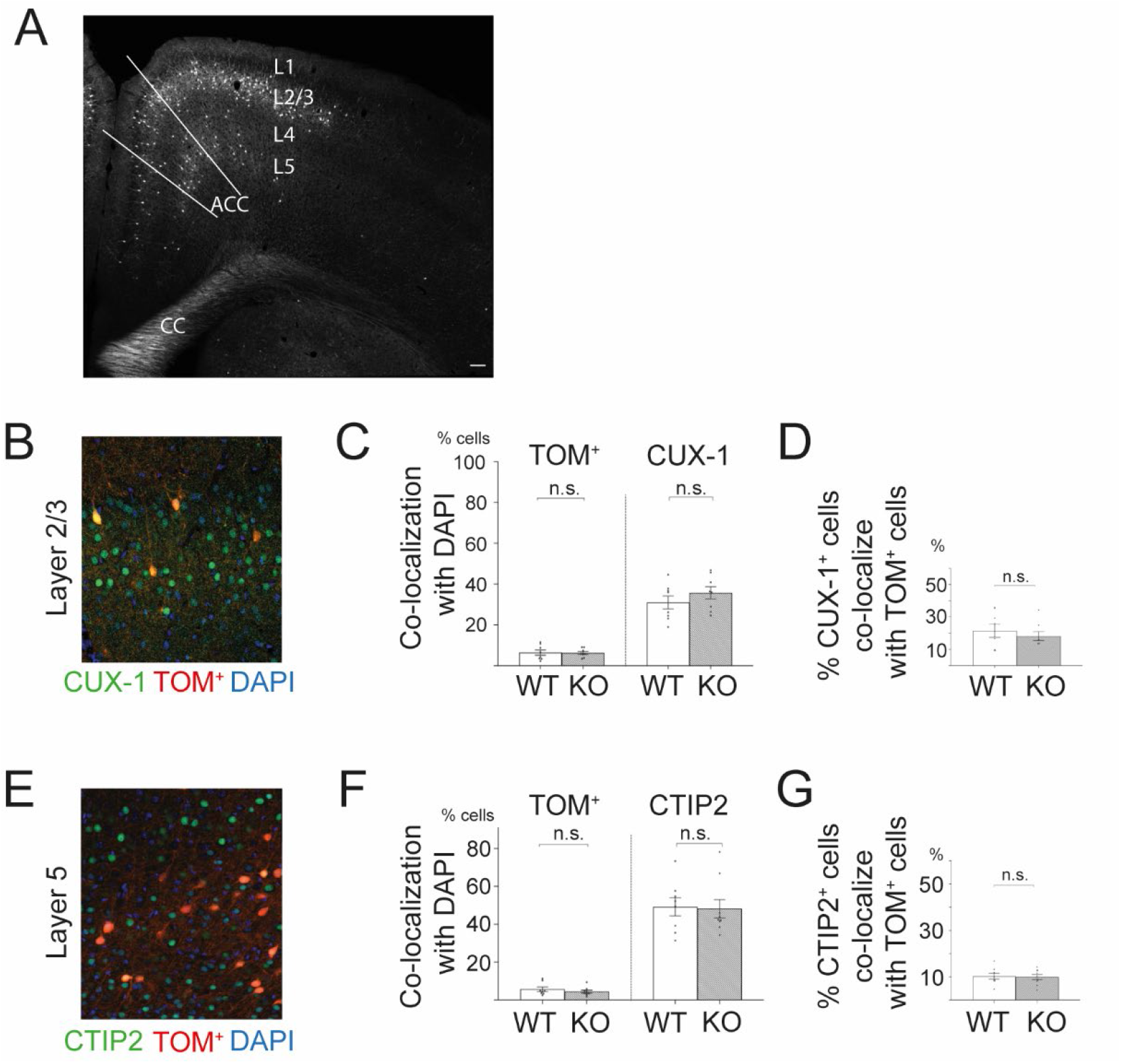
Similar percentages of tdTomato-positive projection neurons are found in L2/3 and L5 of the ACC of WT and *Cntnap2* KO mice. **(A)** Coronal section of the prefrontal cortex from a *Grp:cre, tdTomato^+^*mouse at P14, highlighting the location of the ACC, and the position of TOM^+^ cells in the respective layers. **(B)** A representative coronal section shows staining of the ACC with an anti-CUX-1 antibody (for layer 2/3 projection neurons; green); TOM^+^ cells (red), and DAPI (all cell nuclei, blue). The percentage of TOM^+^ cells, and of CUX-1^+^ cells, which colocalize with DAPI (all cells), is similar between WT and KO. **(D)** The percentage of TOM^+^ cells co-localising with CUX-1^+^ cells is similar between WT and KO mice. **(E)** Layer 5 projection neurons stained for CTIP2 (green), combined with DAPI staining (blue), and TOM^+^ cells (red). **(F)** Percentages of TOM^+^ and CTIP2^+^ cells colocalising with DAPI (all cells) are similar between WT and KO. **(G)** Roughly 9% of CTIP2-positive cells are TOM^+^ cells in both WT and KO. Percentages in charts represent means. Error bars represent S.E.M. WT: n=4; KO: n=4. CC, corpus callosum. Scale bar in (A) = 100 μm.

We analysed dendritic spine densities of pyramidal neurons at four postnatal (P) developmental timepoints: P8, P14, P28 and P56 – that is, approximately, one, two, four and eight weeks of age (Fig. 3). This phase covers the critical time period of circuit formation in this particular area of the cortex (Chini & Hanganu-Opatz 2021, Kroon et al 2019), and allows us to detect transient changes in spine densities. This period also covers the ages for which the behavioural analyses were performed (5 weeks). As with behaviour, we stratified our analyses for male and female *Cntnap2* KO and WT mice.

**Fig. 3.**
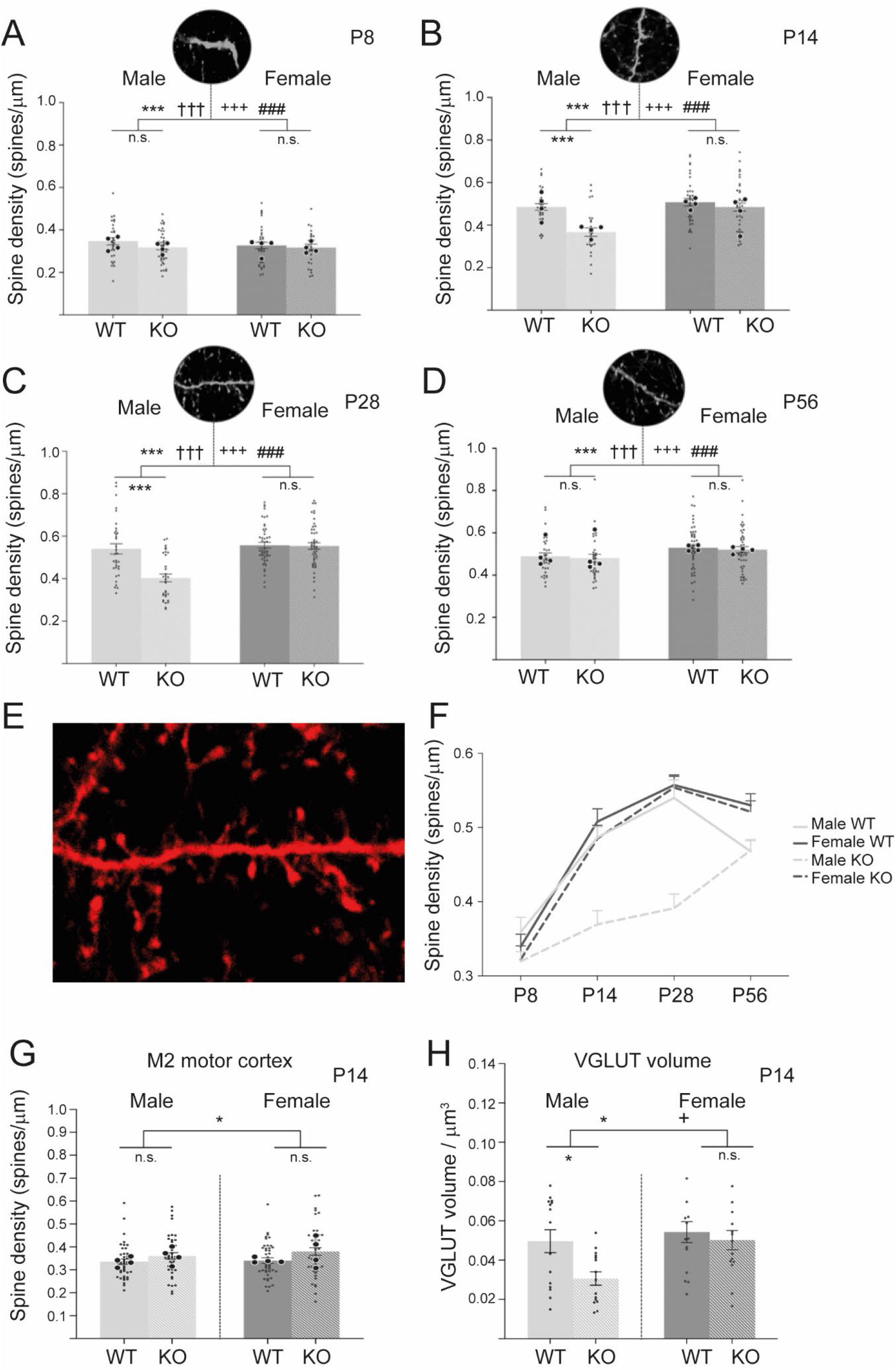
Transient reduction in dendritic spine density in the ACC of male – but not female - *Cntnap2* KO mice. Spine densities in layer 1 of the ACC of *Cntnap2* KO and WT mice from male and female populations were analysed at four developmental time points P8, P14, P28 and P56. **(A)** At P8, no reduction in spine density were observed between WT and KO for both sexes, with male WT (light grey) and male *Cntnap2* KO (striped, light grey), and female WT (grey) and female *Cntnap2* KO (stripped, grey). **(B)** and (C) At P14 and P28, a reduction in spine densities was observed for male KO compared to male WT, and no changes between female KO and female WT. **(D).** At P56, no reduction in spine densities were observed between WT and KO for both sexes. **(E)** Representative images of dendrites and post-synaptic spines made visible using the *Grp-Cre^+^; tdTomato^+^* mouse model which labels a sub-population of pyramidal cells in the ACC (see Fig. 3). White triangles indicate counted spines. a: male WT; b: male KO; c: female WT; d: female KO. **(F)** Overview of the spine densities of WT and KO mice of both sexes at the four different time points. **(G)** No differences in spine densities at P14 in L1 of the secondary motor cortex between WT (grey) and *Cntnap2* KO (striped, light grey) mice of both sexes. **(H)** Density of pre-synaptic boutons (visualized by staining for VGLUT1) at P14 in WT and KO mice of both sexes. A significant reduction in pre-synapse density was observed in male KO vs. male WT mice, but not between female KO and female WT mice. In all experiments, n=4 KO, n= 4 WT for each timepoint and sex. For spine densities, a total of 29 to 48 dendrites were counted for each group. Larger dots represent mean spine density for individual brains, smaller dots the densities of individual dendrites. Statistical analysis for A-D was performed using 3-way ANOVA, although to reduce complexity the data is presented by age. Statistical analysis for G and H was via 2-way ANOVA. Symbols above the bars represent an overall effect of genotype (*), sex (+), age (†) or interaction (#), while symbols below the chart represent Tukey’s *post-hoc* significance between WT and *Cntnap2* KO (*). In each case 1, 2, or 3 symbols represents p < 0.05, p < 0.01, or p < 0.001. Error bars represent S.E.M

Three-way ANOVA revealed a significant effect of genotype, age, sex, and 3-way interaction on spine density in layer 1 of the ACC (F1,576 = genotype: 25.1, p < 0.001; F1,576 = sex: 29.3, p < 0.001; F3,576 = age: 99.8, p < 0.001; F3,576 = 3-way interaction: 3.66, p=0.01). In terms of our investigation into the effect of the *Cntnap2* KO, *post-hoc* Tukey’s correction detected that in males, at the earliest developmental timepoint of P8, spine densities were statistically similar between WT and KO mice (Fig. 3A). However, at P14 (Fig. 3B) and P28 (Fig. 3C), we found a significant reduction in spine densities in *Cntnap2* KO mice compared to WT mice (p < 0.001 for both). Upon approaching adulthood at P56, a difference between the KO and WT groups was no longer apparent (Fig. 3D). This transient reduction in dendritic spine density (Fig. 3F) was not observed at any of four developmental timepoints between female *Cntnap2* KO mice and female WT controls (Fig. 3A-D and 3F).

To analyse the specificity of the spine defects observed in the ACC of male *Cntnap2* KO mice, we also analysed spine density in the secondary motor cortex at P14 (M2, adjacent to the ACC) (Fig. 3G). Two-way ANOVA revealed a significant effect of genotype (F1,166, p=0.02) on spine density in Layer 1 of M2, however this did not survive Tukey’s post-hoc correction and there were no significant differences in spine density between male KO and WT mice, nor between female KO and WT mice. This suggests that knockout of *Cntnap2* in male mice does not cause a global reduction in spine density throughout the brain, but may be region specific.

In addition to tdTOMATO-stained postsynaptic structures, we also investigated pre-synaptic structures at P14 using immunohistochemical staining for the (excitatory) pre-synaptic marker VGLUT 1 – a vesicular glutamate transporter protein (Fig. 3H). Two-way ANOVA revealed a significant effect of genotype and sex on the density of pre-synapses in L1 of the ACC (F1,56 = genotype: 5.42, p = 0.02; F1,56 = sex: 5.95, p = 0.02). Using Tukey’s post-hoc testing, it was shown that male *Cntnap2* KOs displayed a significant reduction in the density of pre-synapses compared to male WT mice (p<0.005), while no significant differences were found between female *Cntnap2* KO and WT mouse populations (Fig. 3H).

All outcomes from pre- and post-synaptic density analyses and significant *post-hoc* corrections are documented in the supplementary index (Table S3).

Overall, this analysis of both pre- and postsynaptic structures indicates a reduced synapse density on apical dendrites of L2/3 and L5 pyramidal projection neurons in the ACC of male *Cntnap2* KO mice, but not of female KOs. This loss in synapse density corresponds to a time point where social deficits of male KO mice were observed (5 weeks of age (Fig. 1)).

### Microglia in male – but not female – *Cntnap2* KO mice show a more activated morphology compared to WT mice

We next sought to identify cell types, molecules and mechanisms which might account for the differences seen in dendritic spine densities between male and female *Cntnap2* KO mice. We analysed two possible scenarios: differential expression of CASPR2 protein between male and female mice, and/or the involvement of sexually dimorphic microglia (Guneykaya et al 2018, Hammond et al 2019, Schwarz et al 2012).

Firstly, for the possible sex-specific bias in expression levels of CASPR2, lysates from the dorsal prefrontal cortex of male and female P28 WT mice were analysed by Western blotting (Figs. 4A and 4B). No significant differences in the expression level of the CASPR2 protein between male and female WT mice were detected (t = 0.34, df = 6). In controls, lysates from male *Cntnap2* KO mice showed no detectable band corresponding to CASPR2 protein, confirming the full knock of this gene in our KO mice (Fig. 4A).

**Fig 4.**
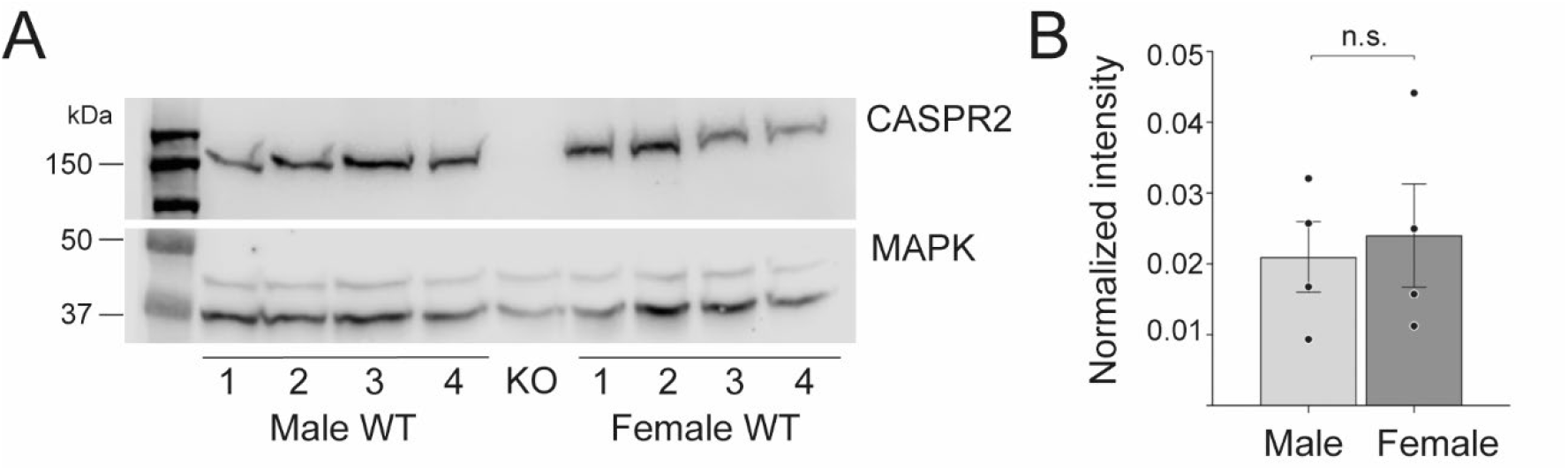
No major differences in expression of Caspr2 between male and female WT mice. **(A)** Western blot analysis of lysates from the dorsal prefrontal cortex (dPFC) of P28 WT mice show no major differences in expression of CASPR2 between male and female (upper part). The expression of MAPK (lower part) was used as a loading control to normalize CASPR2 expression levels. No CASPR2 specific bands were detected in the dPFC from P28 *Cntnap2* KO mice. **(B)** Quantification shows no statistically significant difference in expression levels of CASPR2 protein between male and female WT dPFC brains. Statistical analysis was done using an unpaired 2-tail t-test (t = 0.34, df = 6). Male and Females: n=4 for (labelled 1 – 4 in A). Data are represented as means, error bars represent S.E.M.

Secondly, we investigated the involvement of microglia in the differential effect of the knockout of *Cntnap2* on spine densities in male and female mice. This appeared plausible for a number of reasons. First, microglia are known to play a substantial role in the activity-dependent pruning of synapses in various regions of the brain (Faust et al 2021, Li et al 2019, Paolicelli et al 2011, Schafer et al 2012, Stevens et al 2007), including the ACC (Mallya et al 2019). Microglia are also sexually dimorphic from early postnatal development onwards (Guneykaya et al 2018, Hammond et al 2019, Schwarz et al 2012) and are prominently involved in sculpting the development of neural circuits controlling sex-specific behaviour (McCarthy et al 2017). Besides that, there is evidence that dysfunctional microglia–neuron interactions may play a role in the aetiology of ASD (Li & Barres 2018, Neniskyte & Gross 2017).

To begin with, we analysed the morphology of microglia, a parameter that is closely linked to their function (Vainchtein & Molofsky 2020). Broadly speaking, microglia exist in two contrasting morphological types, resting (called ramified) and activated (called amoeboid), which border the wide spectrum of intermediate phenotypes depending on the strength of their activation (Paolicelli et al 2022). Under physiological conditions, microglia are quiescent/resting, which is morphologically characterised by a small cell body and extensive fine branching processes. These processes are highly motile and constantly survey the local environment. Upon perturbations in the CNS, microglia become activated, undergoing significant changes in morphology, in which the cell body becomes larger and ameboid-like with shorter, thicker pseudopodia which enables microglia to migrate and show higher phagocytic activity (Paolicelli et al 2022).

To discriminate between resting and activated microglia, we analysed their morphology in five different ways – number of primary processes, total number of branch points, total number of branch levels, total process length and total microglia volume (Fig. 5A-5F) – at the two developmental timepoints of P8 and P14.

**Fig. 5.**
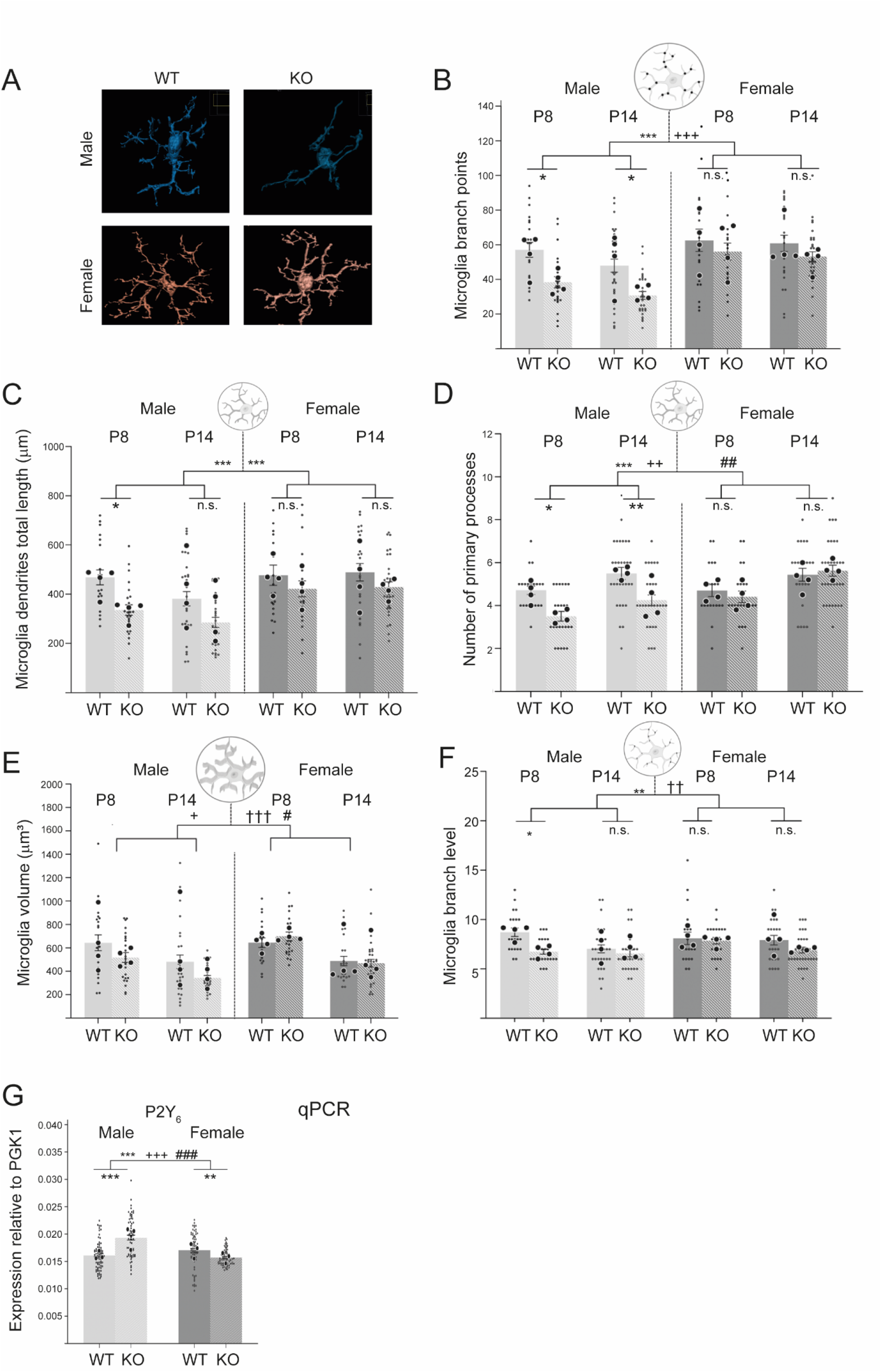
Microglia in male *Cntnap2* KO mice show an altered morphology compared those in male WT mice, with no differences between female WT and KO mice. Microglial morphologies were investigated in layer 1 of the ACC of WT and *Cntnap2* KO male and female mice at different ages. **(A)** Representative images of microglial surfaces reproduced in Imaris. **(B)** Total number of microglial branch points. At both P8 and P14, male KOs had significantly fewer branch points in their processes than WT mice. There were no differences in the female populations. **(C)** Total dendritic length of all processes. At P8, male KOs had significantly reduced total lengths of microglial processes compared to WT mice. There were no differences in the female populations. **(D)** Number of microglial primary processes. At both P8 and P14, male KOs had significantly fewer primary processes than WT mice. There were no differences in the female populations. **(E)** Total microglial volume. There were no differences in the male or female populations. For all microglial morphology experiments, n=4 for KO, and n= 4 for WT, for each timepoint and sex. 3-4 slices were analysed per mouse. Statistical analysis for B-G was performed using 3-way ANOVA. For microglial cell density the data is presented by sex in individual charts to reduce complexity. **(F)** Total number of microglial branch levels. At P8, male KOs had significantly reduced numbers of branch levels compared to WT mice. There were no differences in the female populations. **(G)** qPCR analysis shows an increased expression of the microglial phagocytosis receptor P2Y_6_. In male KO mice compared to male WT mice. Large dots represent results from 3 independent qPCR experiments in which all four conditions (male and female, WT and KO) were analysed in parallel (on the same 96 well plate). Small dot represents individual results from 5 to 6 independent brain preparations of the dPFC of P14 mice (see also Materials and Methods). Statistical analysis for H was performed using 2-way ANOVA. Symbols above the bars represent an overall effect of genotype (*), sex (+), age (†) or interaction (#), while symbols below the chart represent Tukey’s *post-hoc* significance between WT and *Cntnap2* KO (*). In each case 1, 2, or 3 symbols represents p < 0.05, p < 0.01, or p < 0.001. Data presented as means, error bars represent S.E.M. Microglia images from Biorender.

For the mean number of microglia branch points (Fig. 5B), three-way ANOVA revealed a significant effect of genotype and sex (F1,193 = Genotype: 19.44, p < 0.001; Sex: 26.61, p < 0.001). Tukey’s post-hoc analysis indicated that microglia in male *Cntnap2* KOs had a significantly reduced number of branch points at both P8 and P14 compared with microglia in male WT. No differences were found between female *Cntnap2* KOs and WTs at either timepoint (Fig. 5B).

For the mean length of microglia dendrites (Fig. 5C), there was a significant effect of genotype and sex (F1,192 = Genotype: 17.67, p < 0.001; Sex: 17.89, p < 0.001). As with branch level, post-hoc analysis, revealed that the genotype effect was restricted male P8 mice, where *Cntnap2* KOs displayed a significantly smaller mean dendrite length compared with male WTs (Fig. 5C).

For the mean number of primary processes (Fig. 5D), there was a significant effect of genotype, sex and an interaction between genotype and sex (F1,192 = Genotype: 17.67, p < 0.001; Sex: 17.89, p < 0.001; genotype/sex interaction: 9.73, p = 0.002). Post-hoc analysis indicated that microglia in male *Cntnap2* KOs had a significantly reduced number of primary processes at both P8 and P14 compared with microglia in male WT. No differences were found between female *Cntnap2* KOs and WTs at either timepoint (Fig. 5D).

In terms of the volume of the microglia (Fig. 5E), there was no effect of genotype, but there was for age, sex and an interaction between genotype and sex (F1,192 = Sex: 6.176, p = 0.01; F3,192 = Age: 32.67, p < 0.001; genotype/sex interaction: 5.56, p = 0.02).

For the mean number of microglia branch levels (Fig. 5F), a significant effect of genotype, age and 3-way interaction was recorded (F,192 = Genotype: 19.44, p < 0.001; age: 26.61, p < 0.001; 3-way interaction: 4.07, p < 0.05). *Post-hoc* correction revealed that at P8, male Cntnap2 KO mice displayed a significantly reduced number of branch levels compared with male WTs. There was no difference between the two male groups at P14, nor in the two female groups at either timepoint (Fig. 5F).

Overall, in terms of our investigation into the effect of KO of *Cntnap2*, at both P8 and P14, male microglia KOs had significantly fewer primary processes and fewer total branch points, while those at P8 had a significantly smaller total process length and branch levels than microglia from male WT mice (Fig. 5B-5D, 5F). However, the *Cntnap2* KOs were not significantly smaller in volume than the WTs at any timepoint (Fig. 5E). In contrast, there were no differences between female *Cntnap2* KO and WT mice for any of the five morphological measurements at either timepoint (Fig. 5B-5F).

We then investigated whether this reduction in microglia complexity in male *Cntnap2* KOs was also associated with a reduction in the number of microglia within the ACC. We measured microglia cell density in layer 1 of the ACC at 4 developmental timepoints, P8, P14, P28 and P56 (Suppl Fig. 3). Three-way ANOVA revealed a significant interaction between genotype, age and an interaction (F1,460 = Genotype: 26.61, p < 0.001; F3,460 = Age: 26.91, p < 0.001; sex/age interaction: 2.75, p = 0.04). Upon Tukey’s post-hoc correction, it was found that at P8 and P14, there were no significant differences between WT and *Cntnap2* KO mice for both sexes. However, at P28 (p = 0.03) and P56 (p < 0.001), microglia cell density was significantly higher in male *Cntnap2* KO mice compared to male WT mice.

There were no significant differences between female WT and KO mice at any of these four time points (Suppl. Fig. 3).

All outcomes from the microglia analyses and significant *post-hoc* corrections are documented in the supplementary index (Table S4).

### Transcriptional upregulation of a mediator of microglial phagocytosis only in male *Cntnap2* KO mice

To further investigate the differential microglial phagocytosis in male versus female *Cntnap2* mutant mice at P14, we used qPCR to analyse transcriptional changes of the P2Y_6_ receptor – a key regulator of microglial phagocytosis (Fig. 5G). We chose this receptor since it was shown that upregulation of the P2Y_6_ receptor in response to damaged neurons and synapses triggers an increase in microglial phagocytosis (Dundee et al 2023, Koizumi et al 2007). P2Y_6_ is not considered as a chemotactic receptor (i.e. one that responds to a ‘find-me’ signal leading to an attraction of microglia to sites of damaged/non-functional synapses), but as a receptor activated by an ‘eat-me’ signal leading to an increase in microglial phagocytosis (Koizumi et al 2007).

Using qPCR analysis of RNA purified from the dorsal PFC at P14, two-way ANOVA revealed a significant effect of genotype, sex and interaction (F1,276 = Genotype: 8.27, p = 0.004; Sex: 16.45, p < 0.001; Interaction: 47.21, p < 0.001). Tukey’s post-hoc correction showed a transcriptional upregulation of P2Y_6_ in male KO mice compared to male WT mice (p < 0.001), while there was also slight drop in its expression in female KO mice compared to female WT mice (p = 0.03) (Fig. 5G).

This male *Cntnap2* KO specific transcriptional upregulation of a key regulator of microglial phagocytosis, P2Y_6_ is in line with the changes we observed for microglial morphology and synaptic phagocytosis, and further substantiates our finding of an increased microglial synaptic phagocytosis activity only in male KO mice.

In summary, the morphological features of microglia in male *Cntnap2* KO mice compared with those of male WT mice and the qPCR data are therefore indicative of increased phagocytotic (synaptic pruning) activity during early stages of development. Further investigation was therefore required to see if these more active microglia could be involved directly in the reduction in spine density observed for male *Cntnap2* KO mice (Fig. 3).

### Male – but not female – *Cntnap2* KO mice display a greater rate of microglial pre-synapse engulfment than WT mice

Microglia-mediated synapse pruning has been proposed to depend on a sequence of distinct and sequential steps: chemotaxis, target recognition (often including components of the complement system (Schafer et al 2012, Stevens et al 2007), and phagocytosis (Wilton et al 2019). To better understand the meaning of the altered morphology of microglia in male *Cntnap2* KO mice (Fig. 5), and to assess whether microglia directly interact with excitatory synapses during development, we quantified the volume of co-localisation between microglia and pre-synaptic structures (Fig. 6). The analysis was performed at P14, a time point where we had already seen altered microglia morphology as well as differences in spine density between male *Cntnap2* WT and KO mice (Fig. 3).

**Fig. 6.**
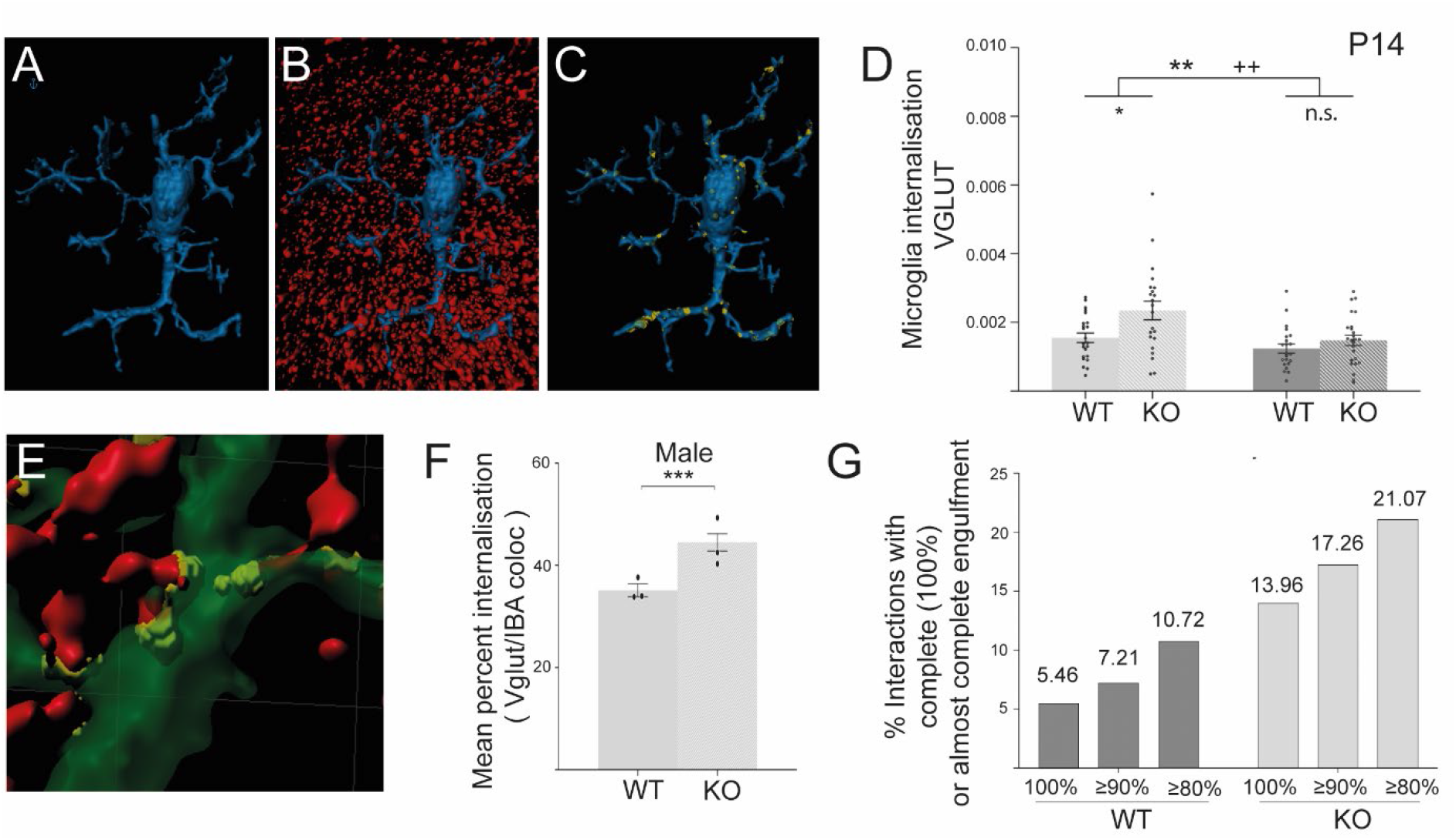
Microglial engulfment of presynapses is more extended in male than female *Cntnap2* KO mice. The co-localisation of microglia (α-IBA stain) and pre-synapses (α-VGLUT1 stain) in layer 1 of the ACC was analysed for P14 WT and *Cntnap2* KO male and female populations. **(A-C)** Representative images of microglia surface rendering (blue)**(A)**, presynapse (red) and microglia surface (blue) rendering **(B)**, and co-localisation between the two (yellow) **(C)**. **(D)** Internalisation of pre-synapses by microglia, normalised to microglia size (volume) and pre-synaptic bouton density (volume). Microglia of male KO mice display a higher internalisation of pre-synapses than microglia of male WT mice. There was no difference for females. **(E)** Representative image of the volume of microglia (green), the volumes of whole pre-synapses (red), and the volumes of their co-localisation, i.e. presumed internalisation (yellow to yellow-greenish), which were used to calculate the percentage internalisation. **(F)** Mean percentage of individual internalised pre-synapses for male mice. Microglia in male KOs display a higher mean percentage internalisation than microglia in male WTs. **(G)** Categorisation of microglia/presynapse interactions based on different levels of engulfment (80% to 100%) shows an overall higher engulfment of presynapses in male KO mice compared to WT. For all experiments, n=4 for KO, and n= 4 for WT, for each timepoint and sex. 3-4 slices were analysed per mouse. Statistical analysis for D was performed using 2-way ANOVA Symbols above the bars represent an overall effect of genotype (*), sex (+), age (†) or interaction (#), while symbols below the chart represent Tukey’s *post-hoc* significance between WT and *Cntnap2* KO (*). Statistical analysis for F was analysed using an unpaired 2-tail t-test. In each case 1, 2, or 3 symbols represents p < 0.05, p < 0.01, or p < 0.001. Data presented as means, error bars represent S.E.M.

We investigated pre-synaptic structures because previous research indicated that microglia target pre-synaptic boutons rather than post-synaptic spines in a process termed trogocytosis (Favuzzi et al 2021, Lim & Ruthazer 2021, Weinhard et al 2018). For a presynapse/microglia co-localisation analysis, we co-immunostained sections from the ACC for VGLUT-1, a vesicular glutamate transporter for (excitatory) presynapses, and for microglia using an α-IBA-1 antibody, in both sexes and genotypes. After normalising for microglia volume and synapse density, two-way ANOVA revealed a significant effect of genotype and sex (F1,88 = Genotype: 8.38, p = 0.005; Sex: 11.07, p < 0.001). Tukey’s post-hoc correction, revealed that at P14 the percentage volume of pre-synapses colocalising with microglia was significantly higher in male KO mice than male WT mice (p < 0.01).

There was no difference between female WT and female *Cntnap2* KO mice (Fig. 6A-6D).

To analyse whether this colocalisation in the males was simply a contact between microglia and pre-synapses or an actual engulfment, we measured what percentage of each presynapse (each VGLUT-1 data point) was internalised by microglia. We found that the mean percentage of internalisation was significantly higher in male KOs than in male WTs, with an average 44.5% of VGLUT-1 being internalised in KO mice versus 35.1% in WT mice (U: 84983, p<0.001) (Fig. 6E-6F).

Defining what constitutes complete engulfment at this resolution is uncertain (Weinhard et al 2018). When setting a threshold of 100% internalisation as a definition for complete engulfment, we found this complete engulfment in 13.96% of all microglia/pre-synapse interactions in KO mice, compared to 5.46% in WT mice (Fig. 6G).

In summary, our data show that at P14, microglia in male *Cntnap2* KO mice display an increased engulfment of excitatory pre-synaptic structures in the ACC relative to controls. Microglia therefore represent a prime candidate for contributing to the reduction in spine densities observed in P14 male KO mice.

## Discussion

### Disruption of social behaviour correlates with decreased spine densities in the ACC of male *Cntnap2* KO mice

*Cntnap2* KO mice represent a well-established genetic model system for autism. Previous behavioural analyses of these mice has revealed deficits in social interaction, communication and stereotypic motor movements. However, to our knowledge none of these studies investigating these mice stratified their investigations by males and females. Our work has now uncovered sex-specific behavioural differences in these mice, with only male KOs exhibiting defects in social interactions, while female KOs remain unaffected (Fig. 1). We did not measure communication or motor stereotypies in our study so cannot comment on whether these behaviours might also show sex differences.

While we did not observe obvious changes in locomotor activity in a 10 min open field test, *Cntnap2* mutant mice have been reported to show a hyperactivity phenotype in the three-chamber social approach test with a complex involvement of the microbiome (Buffington et al 2021) in addition to social deficits. One explanation for the differences seen between these studies may be that different behavioural tests were run, and breeding protocols with Buffington et al. (2021), for example, using offspring from homozygous breeding pairs while we used offspring from heterozygous breeding pairs.

To identify the brain region/s linked to the social deficits, we focussed on the ACC, a prominent and well-studied hub controlling social behaviour whose importance in triggering ASD-like phenotypes has been highlighted in earlier mouse KO studies (Guo et al 2019).

Here we found that reduced spine densities on apical dendrites of L2/L3 and L5 pyramidal excitatory neurons occurred only in male KO mice, while spine densities in female KOs remained similar to those of female WTs (Fig. 3). While not stratified for sex, similar reductions in spine densities of L2/3 pyramidal neurons have been identified in the prelimbic mPFC of *Cntnap2* KO mice (Brumback et al 2018, Lazaro et al 2019), a region close to but distinct from the ACC (Apps et al 2016), but also in cortical layer 5B of the primary somatosensory cortex (Gdalyahu et al 2015). This suggests that dendritic spine reductions in *Cntnap2* KO mice are not confined to the ACC, but reflect the expression of *Cntnap2* in specific regions of the developing cortex (Gordon et al 2016). In addition, we found no reductions in the secondary motor cortex highlighting that knockout of *Cntnap2* in male mice does not cause a global reduction in spine density throughout the (male) brain, but may be region specific.

The decrease in spine density in the ACC we report is in line with a combined rs-fMRI / c-fos study of *Cntnap2* KO mice that demonstrated *hypo*-connectivity between brain regions involved in social interactions, but *hyper*-connectivity between these ‘social’ regions and those not involved in social interactions (Choe et al 2022). These findings fit well with studies in humans and illustrate the complexity of region-specific changes in synapse densities in the brains of ASD individuals (Hutsler & Zhang 2010, Tang et al 2014).

### The decrease in spine densities in male KO mice is transient

Interestingly, the reduction in spine densities in male KO mice is transient and only observed at P14 and P28 of ACC development, while at earlier and later time points (P8 and P56) the spine numbers in male *Cntnap2* KO mice are roughly the same as in WT male mice (Fig. 3). Female *Cntnap2* KO and WT mice do not show statistically significant differences throughout development (Fig. 3). It therefore appears that males have a particular sensitivity to loss of CASPR2 during a critical developmental time window.

CASPR2, the protein product of the *Cntnap2* gene, is a synaptic cell adhesion molecule which has been shown to stabilize already formed synapses, while having little synaptogenic activity (Anderson et al 2012, Gdalyahu et al 2015, Varea et al 2015). Thus, it is tempting to speculate that *in vivo* the initial formation of synapses in male and female KO mice proceeds normally in the first postnatal week, but is followed in the next two to four weeks by a lack of synapse stabilization in *both* male and female KO mice. However, the question then comes up as to why - during the critical period in ACC development - an abnormal number of synapses are removed *only* in male KO but not female KO mice. Our data suggest that the similarity in densities seen as the male mice reach adulthood (P56, 8 weeks (Fig. 3F)) is caused both by a partial recovery in synapses in the mutant mice but also a period of neurotypical synaptic pruning in the male wild types.

It also remains to be analysed in further detail whether this faux recovery in adult male *Cntnap2* KOs that sees a return to spine densities similar to WTs is associated with a return to normal social interactions or whether the social defects remain as it has been observed in fact in other investigations (Scott et al 2019). This transient reduction fits well with the interesting concept that neuro-developmental miswiring during critical stages of early neural circuit development often results in long-term consequences of major functional and behavioural defects that remain in adults (Del Pino et al 2018, Marin 2016).

### Microglia as candidates for the sex-specific differences in spine densities

Why loss of *Cntnap2* should have differential effects on pyramidal synapse development in the two sexes was not immediately clear. The most obvious hypothesis was that there may be a differential expression of the CASPR2 protein in the cortex of male versus female mice. Our findings indicate that this is not the case and that the expression of CASPR2 protein is similar between the two sexes in the dorsal prefrontal cortex (Fig. 4). Previous studies have shown that CASPR2 expression is broadly distributed in the rodent brain, although such results did not differentiate between sexes (Abrahams et al 2007, Gordon et al 2016).

Since microglia are major players in activity-dependent synaptic pruning (Paolicelli et al 2011) and are sexually dimorphic from early postnatal development onwards (Guneykaya et al 2018, Hammond et al 2019, Schwarz et al 2012, Thion et al 2018), they therefore appeared to be a good candidate for the reduced spine densities on dendrites of L2/3 and L5 pyramidal neurons in the ACC. In fact, microglia in male *Cntnap2* KO mice displayed an increased activation status, as well as engulfment and phagocytosis of pre-synaptic structures, compared to male WT mice (Figs. 5 and 6). Furthermore, qPCR showed a transcriptional upregulation of the key microglial phagocytosis receptor P2Y_6_ in male KO mice only (Fig. 5G) while this receptor was even slightly downregulated in female KO mice compared to female WT mice. This provides initial evidence that microglia play a role in mediating the sex-specific deficits in spine densities in the ACC of male *Cntnap2* KO mice through their increased phagocytotic activity.

To our knowledge, this is the first time microglial differences have been reported in the *Cntnap2* KO mouse model and may initially appear surprising. Despite some studies proposing that CASPR2 may be expressed in glial cells (Poliak et al 1999), no study has shown the protein to be expressed in microglia. The only other investigation into microglia in relation to loss of CASPR2 measured their cell density and size, but only in adult KO male mice, finding no differences compared with WTs in the hippocampus, mPFC, and striatum (Cope et al 2016). There are, though, other reports of differences in another glial cell type, the oligodendrocytes, during development. In this case, myelination was found to be delayed in the somatosensory cortex of male *Cntnap2* mutant mice, due to an early deficit in the number of oligodendrocytes (at P21) that was later compensated for in the adult cortex (P56) (Scott et al 2019) – a situation that is strikingly similar to our observation with recovery of post-synaptic spine densities to wild type levels. Taken together these results further stress the importance of both timing and sex when investigating the *Cntnap2* mouse model. There appears to be a key developmental window in these mice, in which the role of *Cntnap2* is particularly important and requires further research in this model.

### Divergent characteristics of male and female microglia

Likely candidates for instigating differences between male and female microglia are sex hormones, in particular testosterone that shows a perinatal surge in male mice only (McCarthy et al 2017). Prior to this transient testosterone increase, there are no differences between the sexes in the number or morphology of microglia in any brain region analysed, however, differences are found in the first postnatal week (Dubbelaar et al 2018, Hanamsagar et al 2017, Nelson et al 2017, Schwarz et al 2012).

Testosterone-induced differences in microglia characteristics play a crucial role later during puberty in refining neural circuits controlling sex specific behaviours (Kopec et al 2018, Lenz et al 2013, VanRyzin et al 2019), however, our data suggest that these differences come – accidently – much earlier into play in response to the abnormal synapse development in *Cntnap2* KO mice. Such an idea would fit into the *errant gardener hypothesis* (Neniskyte & Gross 2017), stating that male (but not female) microglia might show an errant over-reaction in response to destabilized synapses, triggering an over-pruning of synapses, and causing irreversible damage to developing neural circuits, which ultimately leads to behavioural defects in adult mice. Interestingly, there is already evidence that raised levels of 17β-oestradiol can rescue phenotypes in *Cntnap2b* zebrafish mutants. A study of these mutants displayed reductions in inhibitory neurons and dysregulated signalling in excitatory and inhibitory neurons, leading to hyperactivity, a common symptom in ASD. This phenotype was reversed using oestrogen receptor agonists (Hoffman et al 2016).

Our data fit well with recent findings that the cause for the male/female imbalance in ASD is not due to a hypothetical sex-specific expression of ASD risk genes, but that ASD risk genes interact with characteristic dimorphic pathways (Werling et al 2016). In other words, it is not the sex-specific differential expression of ASD candidate genes but rather ‘naturally occurring’ sexually dimorphic processes (here the sexual dimorphism of microglia), that modulate the impact of risk variants (here *Cntnap2* KO) and contribute to the sex-imbalance in ASD.

### Limitations of the study

In terms of behavioural deficits in the *Cntnap2* mouse model, we have investigated here only one aspect, social interactions, but not other ASD-like characteristics such as repetitive behaviour, stereotypic motor movements or ultrasonic vocalisation (Buffington et al 2021, Penagarikano et al 2011, Scott et al 2019). Further experiments will be directed towards an understanding of whether these aspects also show a sexual dimorphism in the *Cntnap2* KO mouse model.

Given microglia are highly reactive to their environment, care had to be taken to control for confounding factors, such as the effect of animal housing, maternal care and microbiome (food). However, it cannot be entirely excluded that variations exist owing to the linear nature of the study.

Analysis of the microglia-pre-synapse colocalisation is limited by the resolution of the Zeiss confocal microscope and 60x lens used. It is likely that any colocalisation is at the very limits of the resolution, making it likely that any colocalisation is of groups of pre-synaptic boutons rather than individual ones. Despite this being a commonly used technique (Nemes-Baran & DeSilva 2021), confirmation via electron microscopy and/or analysis with a marker for phagocytosis (e.g., CD68) is therefore required.

Given that our data are derived from one particular mouse model of ASD they can only provide some rather restricted concepts towards the aetiology of ASD in humans. Our results need to be further investigated in other principle ASD mouse models (Ellegood et al 2015) to better understand the different aetiologies in autism as a spectrum disorder (Buch et al 2023).

## Materials and methods

### Mouse work

This study was performed in strict accordance with UK and EU regulations, with approval granted by the King’s College London Ethics Committee. All mice were bred on site at the King’s College London Biological Services Unit (BSU) on Guy’s Campus. All procedures were performed with UK Home Office project and personal licenses in accordance with the Animals (Scientific Procedures) Act 1986. Unless stated otherwise, mice were euthanised in a chamber of rising carbon dioxide, with death confirmed by cervical dislocation. In order to adhere with King’s College London guidance on the 3Rs, three or four animals were used for any reported quantitative analysis when power analysis allowed. Number of animals used for each experiment are detailed in Figure legends.

### Mouse lines

B6.129(Cg)-*Cntnap2^tm1Pele^*/J mice carrying loss-of-function *Cntnap2* alleles were obtained from O. Marin (CDN, King’s College London)(Scott et al 2019) and originally derived from the Jax Lab (Bar Harbor, Maine, USA). These mice were backcrossed into a C57BL/6 J background for >10 generations. *Grp-Cre_KH288* mice (Gerfen et al 2013) were bred with the B6.Cg-Gt(ROSA)26Sor^tm14(CAG-tdTomato)Hze^/J mouse line (Jax Lab) that contain in the Rosa26 locus a stop-floxed cassette for expression of the fluorescent protein TdTomato (TOM), so that TdTomato is expressed in a Cre-dependent manner. In *Grp-Cre_KH288* mice, Cre expression is mostly limited to pyramidal cells of L2/3 and of L5 of the prefrontal cortex (Fig. 3). Onset of CRE expression was observed during the first postnatal week. Due to the local expression of CRE (determined by the *Grp* promoter), tdTomato expression in *Grp-cre^+^; tdTomato^Flox^* mice is mostly confined to L2/3 and L5 in the PFC. These mice were backcrossed into a C57BL/6 J background for >10 generations.

*Grp-Cre^+^; tdTomato^+^* mice were bred for >10 generations with *Cntnap2* heterozygous mice (*Cntnap2* ^+/-^). Heterozygous offspring were used for breeding to generate *Grp-Cre^+^; tdTomato^+^; Cntnap2^-/-^* (hereafter called *Cntnap2* KO) and *Grp-Cre^+^; tdTomato^+^; Cntnap2^+/+^* (hereafter called WT) littermates, which were used in all experiments. *Cntnap2* ^+/-^ mice were not analysed in this study. To obtain time-mated females, male and female mice were placed together overnight and if a vaginal plug was detected the following day it was designated P0.

*Cntnap2* genotypes were determined by PCR using primers recommended by the Jax lab, with CASPR2.3 Neo *TTG GGT GGA GAG GCT ATT CGG CTA TG;* CASPR2.4 *AAG AGG GTG AGG GGA GAA AA; and* CASPR2.5 *CAA GAA TTG GGA ATG CAC CT*.

### Sex determination

All P8 mice were sex determined twice, by visual inspection by the experimenter, and by using specific PCR primer sets using protocols described in Tunster et al. (2017).

### Mouse husbandry for behavioural experiments

All mice were housed in Techniplast cages (32 cm x 16 cm x 14 cm) with sawdust (Litaspen premium, Datesand Ltd, Manchester, UK) and basic cage enrichment, consisting of sizzlenest (Datesand Ltd, Manchester, UK) and a cardboard shelter (LBS Biotech, Horley, UK). Cages were never cleaned the day before, or on the day of testing in order to minimise the potential effects of cage disturbance on the behaviour of the mice. All mice had *ad libitum* access to water and food (PicoLab® Irradiated Rodent Diet 205053, LabDiet, USA). The housing room was maintained at constant room temperature (∼21 °C) and humidity (∼45%) and kept under a regular light/dark schedule with lights on from 08:00 to 20:00 hours (light = 270 lux). Test mice were housed matched by sex (2-3 mice/cage) when weaned at 4 weeks of age. This has previously been shown to have minimal effects on C57BL/6J mice and eliminates the potential confounds of large group housing, such as the establishment of social hierarchies (Brown 1953, Lad et al 2010). The oestrous phase of the female mice was not checked in this study, but it is unlikely that this affected the results as there were no major differences in the variance observed in the behavioural measures between males and females (Beery 2018). For the behavioural experiments, littermate controls were used, and test mice were obtained from 11 different litters to avoid litter-specific effects on behaviour (Jimenez & Zylka 2021). C57BL/6J male and female conspecifics for social tests (social investigation) were purchased from Charles River (Margate, UK) one week before testing to allow for a habituation period. These conspecific mice were always pair-housed by sex and kept in different holding rooms to prevent any exposure to the test animals before social testing.

### Behavioural testing

All behavioural tests were performed in the light phase between 09:00 and 18:00 hours. Although mice are nocturnal animals, previous research suggests that many behaviours are not affected by the time of the day mice are tested, but by the change in light levels, from housing to testing environments (Beeler et al 2006, Valentinuzzi et al 2000). For all experiments and behavioural scoring, the experimenter was blind to the genotype.

Behavioural testing began when mice were 4 weeks old, when mice are still considered juvenile. The first three tests were open field (Candland & Nagy 1969), social interaction (unfamiliar test mice matched by age, sex and genotype), and social investigation (with an unfamiliar and sex-matched juvenile C57BL/6J mouse (Grayton et al 2013, McFarlane et al 2008). From 8-12 weeks of age, mice were tested as adults: open field (Candland & Nagy 1969); social investigation (Yang & Crawley 2009). A social interaction test between adult paired test mice was not conducted for welfare reasons due to risk of aggression between unfamiliar adult mice.

At least one day intertrial interval was included between different tests. All tests were recorded using a camera positioned above the test arenas and movement of each mouse tracked using EthoVision software. After each trial, boli and urine were removed from the test arena which was then cleaned with 1 % Anistel® solution (Tristel Solutions Ltd, UK). At the end of testing, mice were returned to their home cage, which was returned to the housing room. Light levels for each task varied according to the specific task.

### Open field test

The open field test was carried out as adapted from Candland and Nagy (1969). Individual mice were placed in a circular arena with a diameter of 40 cm for 10 min, with light level at the centre of the arena at 10 lux (white light). A 20 cm diameter circle at the centre of the open field arena was designated the centre zone, a ring 20 cm thick around the perimeter of the arena designated as the outer zone. Avoidance of the centre zone was used as a measure of anxiety, and total distance moved in the outer zone was used as a measure of locomotive activity.

### Social interaction (unfamiliar test mice matched by sex and genotype)

Social interaction between paired juvenile test mice was assessed as described above in the social investigation test with a few modifications. Test mice unfamiliar with each other were paired, matched by sex and genotype (i.e., male wildtype pairs, male *Cntnap2*^-/-^ pairs, female wildtype pairs, female *Cntnap2*^-/-^ pairs). Members of each test pair were individually transferred to the test room and placed together in a clean cage, identical to their normal home cage (Techniplast cage, 32 cm x 16 cm x 14 cm) containing only sawdust at a height of 2 cm. The testing area was dimly lit from below (10 lux white light). The tails of the mice were uniquely marked within each pair with a non-toxic marker pen (Pentel, UK) prior to testing so they could be identified. Interactions initiated by either member of the mouse pair were recorded for 10 min and social behaviours were later scored from the recordings by researchers who were blind to the mouse genotypes and sex. If aggression is observed for prolonged periods (longer than 30 s in one continuous bout or >2 min total cumulative duration), the trial is stopped, and the pairs separated. In this study, no trials exceeded the threshold for terminating the trials early. Following testing, the mice were returned to their respective home cages (Terranova & Laviola 2005, Winslow 2003).

### Social investigation with an unfamiliar, sex-matched C57BL/6J mouse

Social investigations were assessed as described previously (Grayton et al 2013, McFarlane et al 2008). Prior to testing, mice were transferred to a clean cage, identical to their normal home cage (Techniplast cage, 32 cm x 16 cm x 14 cm) containing only sawdust at a height of 2 cm and habituated to this environment for 1 hour. The testing area was dimly lit from below (10 lux white light). The tails of the conspecific mice were marked with a non-toxic marker pen (Pentel, UK) so they could be identified in the recording. During testing, mice were transferred into the testing room in their ‘new’ home cage, and a sex-matched novel conspecific mouse (C57BL/6J aged 4 weeks) was put into the cage with them. Investigations initiated by the test mouse were recorded for 10 min and social behaviours were later scored from the recordings by researchers who were blind to the mouse genotypes. If aggression was observed for prolonged periods (longer than 30 s in one continuous bout or >2 min total cumulative duration), the trial was stopped, and the pairs separated. In this study, three trials with male mice (one trial with WT and two trials with KO) exceeded the threshold for terminating the trials early. Following testing, the mice were returned to their respective home cages (Terranova & Laviola 2005, Winslow 2003). The number of social sniffing (sniffing above the middle of the body), anogenital sniffing (sniffing below the middle of the body) and bouts of following (mouse moves near the other mouse, follows its movements without making direct contact with the mouse) behaviours made by either test mouse towards the other test mouse were recorded (Terranova & Laviola 2005, Winslow 2003).

### Olfactory habituation/dishabituation

Olfactory habituation/dishabituation is used to assess deficits in olfaction (Yang & Crawley 2009). As social behaviour relies heavily on olfaction (Zou et al 2015), this test is a necessary control for the interpretation of the social behaviour. Animals were tested in their home cage, which had been cleaned out 2-3 days prior to testing, with all enrichment removed and a fresh cage lid to minimise interfering odours. Following a 10 min habituation the mouse was exposed to three odours in turn: water (control/no odour; 50 μl), vanilla essence (non-social; 50 μl, 1:100 dilution; Uncle Roy’s, Moffat, UK) and urine collected from a unfamiliar, sex-matched conspecific (social, 25 μl). Each odour was presented on a cotton-tipped wooden applicator 3 times and for a period of 2 min each time with an interval of roughly one min while the next cotton bud was prepared. The total number of sniffs of the cotton bud made by the mouse during every trial was recorded. Habituation to an odour was defined as a decrease in sniffing over consecutive presentation of the same odour, and dishabituation as an increase of sniffing when a new odour is presented. The testing area was dimly lit from below (10 lux white light).

### Antibodies

The following primary antibodies were used: rat anti-CTIP2 1:500 (Abcam ab18465), rabbit anti-CUX1 1:500 (Abcam, ab140042) rabbit anti-IBA-1 primary antibody (Abcam, ab178846), rabbit anti-IBA-1 primary antibody 1:500 (Wako, 4987481428584), rabbit anti-Caspr2 1:1000 (Abcam ab137052), rabbit MAPK1/ERK antibodies 1:1000 (Cell Signalling mAb#4695), rabbit anti-DSred primary antibody 1:500 (Takara, 632496) which binds Td-tomato protein, and guinea pig anti-VGLUT1 1:1000 (Synaptic Systems 135 304).

The following secondary antibodies were used: goat anti-rabbit Alexa-488 secondary antibody 1:1000 (Abcam, ab150089), donkey anti-rabbit Alexa-488 1:1000 (Abcam ab150073), goat anti-guinea pig Alexa-647 secondary antibody 1:1000 (Abcam ab150187), donkey anti-rabbit Alexa-647 1:1000 (ABCAM ab150075), goat anti-rat Alexa-488 1:1000, (Abcam ab150157).

### Histology

Coronal slices were prepared from *Cntnap2* KO and WT littermates aged P8, P14, P28, and P56. For this, postnatal mice were anaesthetised via intraperitoneal injection of 0.1 ml of Pentobarbital and then perfused with 20 ml of ice-cold 1x PBS until the liver had drained of blood, followed by fixation with 70 ml of ice-cold 4% PFA/PBS. Successful perfusion was confirmed by tail and/or muscle twitch, hardened body and a complete white brain. Following perfusion, the brain was extracted and fixed for 2 hours in 4% PFA at 4 °C. Brains were then washed 3 times in 1x PBS.

For sectioning, brains were embedded in 6% agarose/PBS and sectioned coronally at 50 µm using a Leica VT1000S vibratome (P28 and P56) or a Camden 5100mz vibratome (P8 and P14). If storage of sections was required at any point in this process, this was done in PBS, 0.25% Tween (PBST) + Sodium Azide at 4 °C. When sections were selected for staining and imaging, they were matched across brains to keep regions consistent.

### Immunohistochemistry

Free-floating coronal sections were processed for immunohistochemistry using protocols that varied depending upon the primary antibody used. Sections were always placed on a shaker during washing or immune-labelling steps. A list of primary and secondary antibodies with dilutions used are provided under Materials.

For the anti-IBA-1, anti-Ctip2 and anti-DSred primary antibodies, sections were first permeabilised and washed 4 times in 1x PBS, 0.25% Triton (PBST) for 15 minutes each at room temperature (RT). Sections were blocked with 0.1% Triton, 5% BSA, 5% normal goat serum (NGS), 0.2 M glycine in 1x PBS for 2 hours at RT. The primary antibody was then diluted in blocking buffer and sections were incubated in blocking buffer overnight at 4 °C. The following day, sections were then washed in 1x PBS 4 times for 15 minutes each at RT. The secondary antibody was diluted in 1x PBS, 0.1% Triton, 5% BSA, 5% NGS and incubated with the sections for 2 hours at RT. Sections were then washed in 1x PBS 4 times for 15 minutes each at RT.

For the anti-VGLUT1/anti-IBA-1 primary antibody co-staining, sections first underwent antigen retrieval in sodium citrate buffer containing 10 mM Sodium Citrate, 0.05% Tween 20, at pH 6.0, for 30 minutes at 90 °C. Sections were then washed in 1x PBS plus 0.05% Tween20 twice for 2 minutes each, before being permeabilised in 0.5% PBST for 30 minutes at RT. Sections were blocked with 1% Triton, 5% BSA, 5% NGS and 0.2 M glycine in 1x PBS for 3 hours at RT. The primary antibody was then diluted in blocking buffer and sections were incubated for 2 days at 4 °C. After standard washing, the secondary antibody was diluted in blocking buffer and incubated with sections for 2 hours at RT. Sections were then washed in the blocking buffer 3 times for 30 minutes each at room temperature.

Subsequently, all sections, including those with antibody staining, were stained with 4’6-diamidino-2-phenylindole (DAPI) for 5 minutes, before washing twice in 1x PBS and being mounted using Mowiol. Slides were left to dry for 24 hours while protected from light.

### Region identification

To keep consistency between the sections to be analysed, and to aid location of the ACC, 4 sections were mounted from each brain that corresponded most closely to defined regions in The Atlas of the Mouse Brain in Stereotaxic Coordinates (Paxinos & Franklin, 2004) and/or the Allen Mouse Brain Atlas (http://mouse.brain-map.org/). Sections were kept consistent within an experiment but varied between experiments. The endogenous Td-TOMATO fluorescence (in L2/3 and L5) and DAPI staining were used to aid identifying the cortical layers.

### Spine density and morphology measurement and analysis

Male and female WT and KO littermates were selected at the developmental timepoints indicated, and tissue prepared as described above.

Given the experimental mice contained an endogenous tdTOMATO fluorescence, no immunohistochemistry was required except for P8 mice to identify dendrites and spines. In P8 mice, imaging spines using the endogenous tdTOMATO required enhancement using a Rabbit anti-DSred primary antibody which binds also Td-tomato protein.

The *Grp:cre; tdTomato^+^* mouse line sparsely labels pyramidal neurons including their apical dendritic projections to layer 1 and spines (see Fig. 3). From each of four sections per brain, 5-6 dendrites no shorter than 50 µm were imaged in layer 1 of the ACC, giving a total of 15-18 dendrites per brain. Images were acquired with a Zeiss Axio Imager.Z2 LSM 800 Confocal microscope using a 60x oil immersion objective. All dendrites were imaged as a z-series (with optical sections at 0.28-0.30 µm step size), averaged 16 times, with 1024×1024 pixel resolution.

All files were blinded to sex and genotype. Dendritic spines were then manually counted from the z-series images in Fiji (Schindelin et al 2012) using its Cell Counter plugin. Dendritic spine density was calculated by dividing the total number of spines over a given length of dendrite (spines/μm).

### Microglia cell density measurement and analysis

Immunohistochemistry was performed with a rabbit anti-IBA-1 primary antibody, a marker for microglia. For four sections per brain, using a Zeiss Axio Imager.Z2 LSM 800 Confocal microscope and a 20x oil immersion objective, tiled images of the entire medial prefrontal cortex and corpus callosum were collected and analysed.

Images were quantified using FIJI, with all files blinded to sex and genotype. The background was subtracted to reduce noise, and brightness and contrast were adjusted. The Allen Brain Atlas was used as an aid to identify the ACC and mark its region and layers on each of the images using the Freehand tool and Polygon selections. DAPI and the endogenous tdTomato expression were used to delineate and mark the cortical layers within the ACC.

For analysis of IBA-1^+^ microglia density, thresholding was set to a range that captured the entire cell but without adding noise or outliers. This range was then kept consistent throughout the analysis with size threshold limits set to 25–120 µm^2^. Microglia density per mm^2^ of the ACC was then counted manually, using the Analyze Particles tool to confirm results. Soma area and the areas of the cortical layers within the ACC were measured using the Freehand tool.

### Microglia morphology measurement and analysis

For microglia morphology measurements, rabbit anti-IBA-1 primary antibody (Wako) was used showing improved clarity of microglial processes. Microglia morphology was quantified using Imaris 9.1 software (Bitplain, Zurich, Switzerland) but only for microglia whose processes were all within the field of view. Each image was background and baseline corrected. 3D reconstruction of microglia processes was performed using the Imaris FilamentTracer tool by creating a region of interest (ROI) around the microglia. Spot Detection mode helped to determine start and end points, with no loops allowed or points within the cell body. Initial analysis was performed automatically using the software, with manual verification and modification then applied. Three morphometric parameters were analysed: number of primary processes, total process length, and total branch points (Fig. 5). A primary process was defined as an extension projecting from a microglia cell body with a length greater than 3 μm and a width greater than 0.5 μm. Total microglia volume was calculated via the Imaris Surface Tool (Fig.5);(Nemes-Baran & DeSilva 2021).

Absolute intensity threshold settings were determined in a small representative pilot sample first, and then kept consistent thereafter using a range of *1.01e4 and 1.40e4* to allow for individual image background variability. Automatic thresholding was used as an initial guide to find the surface edge of each microglia, being careful not to over- or under-represent the boundary. Any disconnected surfaces were reconnected manually using the Unify feature. Total microglia volume and surface area could then be extracted. Surface renders were then compared to FilamentTracer renders to ensure both techniques accurately represented the microglia morphology.

### VGLUT-1-microglia colocalisation measurement and analysis

Immunohistochemistry was performed via double staining. Microglia were stained using a rabbit anti-IBA-1 primary antibody, pre-synapses of excitatory synapses were stained using a Guinea-pig anti-VGLUT-1 primary antibody. From three sections per brain, 7-8 microglia were imaged in layer 1 of the ACC, giving a total of 21-24 microglia in total.

Co-localization was performed on Imaris using the Surface Tool and Surface-Surface Coloc XTension as described in (Nemes-Baran & DeSilva 2021). Microglia and VGLUT surface renderings were performed as described for the morphology analysis, with the exception that Split Touching Objects was enabled for the VGLUT stain to ensure pre-synapses in close proximity were separated. The volume of VGLUT internalised by microglia was then determined by the Surface-Surface Coloc XTension with no smoothing. This created a new 3D surface render of just colocalised surfaces, from the edge of the microglia inwards. Results were then normalized to the total VGLUT volume within the tile and the total microglia volume.

### Statistical analysis

Power calculations performed using in G∗power (v3.1.9.4). (F-test, ANOVA, fixed effects; power = 0.80, alpha = 0.05) revealed that a group size of n = 11 per group for behavioural analysis were required to achieve p < 0.05, and a group size of n = 4 per group for the spine and microglia analyses were required to achieve p < 0.05. All statistical analyses were performed using Prism (v9.1.0, GraphPad Software Inc., La Jolla, CA, United States). P-values below 0.05 were considered statistically significant. Normality and variance tests were first applied to all experimental data. Behavioural data were analysed using two-way ANOVA with main factors of genotype (WT or KO) and sex (male or female) followed by Tukey’s multiple comparisons test between genotype groups within each sex. Comparison between genotype (WT or KO) and sex (male or female) at multiple developmental timepoints for the spine and microglia analysis were performed with a three-way ANOVA where the data was parametric, or Kruskal-Wallis test where it was non-parametric. Analyses over a single timepoint were performed using two-way ANOVA. *Post-hoc* correction for multiple comparison was performed using the *post-hoc* Tukey’s method (with ANOVA) and Dunn’s method (with Kruskal-Wallis). Comparison between groups was performed using an unpaired two-tailed *t*-test.

### Western blot analysis

Western blot analyses from forebrain lysates were performed using standard techniques. In brief, brains were perfused with ice-cold 1x PBS. Then tissue was isolated from the dorsal PFC of P14 mice using a CytoVista mouse brain slicer (1 mm), isolating for each brain the section optimally including the ACC, and lysed in RIPA buffer including protease inhibitors, denatured at 95 °C for 5 min, spun down, and supernatants were stored in aliquots at -80 °C. After 10% SDS PAGE, proteins were transfer to nitrocellulose filters, blocked in 1% milk powder in TBST (blocking buffer) for 2 h at RT, incubated with mouse anti-CASPR2 antibody (1:1000) overnight at 4 °C, washed 4 times 15 min in TBST, incubated with anti-mouse HRP-coupled secondary antibody (1:5,000) for 2 h at RT in blocking buffer, and washed 4 times 15 min in TBST. Chemiluminescence intensity reflecting protein expression levels were determined according to standard protocols. In parallel, MAPK protein expression was determined for normalisation of the amount of protein loaded. Densitometric analysis was performed using FIJI.

### qPCR analysis

Tissue was isolated from the dorsal PFC of P14 mice using a CytoVista mouse brain slicer, isolating the section optimally including the ACC. Five to six independent brain preparations were performed per condition (WT and KO for both sexes). RNA was isolated using Monarch RNA purification kit, and cDNA using ThermoFisher SuperScript IV cDNA synthesis kit with random 10mers. LightCycler PCR was performed using PCRBiosystems kits. For normalisation, qPCR was performed on the housekeeping gene PGK1 using upstream primer 5’-AGGGTGGACTTCAACGTTCC-3’ and downstream primer 5’-AACGGACTTGGCTCCATTGT-3’ with a product length of 111 bp, and for the P2Y_6_ receptor, upstream primer 5’-CCAAATCTGGCACTTCCTCCT-3’ and downstream primer 5’-TGGATGGTGCCATTGTCCTG-3’, with product length of 117 bp. In both cases, primers were located on different exons separately by large introns. qPCR and its analysis was carried out using a Roche LightCycler96, following instructions by the manufacturer. Each RNA was analysed with four technical replicates in each of three independent experiments (shown as small dots in Fig. 5). In each independent experiment, 5-6 cDNA preps from each of the four conditions were analysed for the expression of P2Y_6_, relative to the house keeping gene PGK1, in parallel on one 96-well plate. Large dots in Fig. 5F show the results of each of these three independent experiments.

## Acknowledgements.

We would like to thank staff from the BSUs at NHH and Denmark Hill, in particular Tiffany Jarvis, for meticulous support of our work, O. Marin (KCL) for providing the *Cntnap2* KO mouse line, Yaroslav Kainov and Anna Zhuravskaya for advice in qPCR technology. We acknowledge valuable comments on the manuscript by S. Guthrie and J. Clarke. This work was supported by a BBSRC grant to U.D. (BB/T013753/1), and a Wellcome Trust Investigator Award to M.C. (103759/Z/14/Z).

## Author contributions

All experiments apart from behavioural experiments were performed by M.D., K.G.-F., and V.T, with L.Y., H.H., and K.M. contributing to the experiments. The behavioural experiments were performed by S.N., K. G.-F., Z.A., and C.F.

The study was designed by U.D., M.D. and C.F. The paper was written by U.D. and M.D., with significant contribution from C.F. and M.C. U.D. supervised all experiments. Final approval was made by U.D.

All authors approved the final version of the manuscript and agreed to be accountable for all aspects of the work. All authors designated as authors qualify for authorship, and all those who qualify for authorship are listed.

## Corresponding author

Correspondence to Uwe Drescher

## Ethical Declarations

Competing interests.

The authors declare no competing interests.

## Ethical approval

This study was performed in strict accordance with UK and EU regulations, with approval granted by the King’s College London Ethics Committee. All mice were bred on site at the King’s College London Biological Services Unit (BSU) on Guy’s Campus. All procedures were performed with UK Home Office project and personal licenses in accordance with the Animals (Scientific Procedures) Act 1986.

## Suppl. Information

See separate file

