## Supplementary material for "Sexual dimorphism in the social behaviour of *Cntnap2* KO mice correlates with disrupted synaptic connectivity and increased microglial activity in the anterior cingulate cortex of males": Suppl Figures

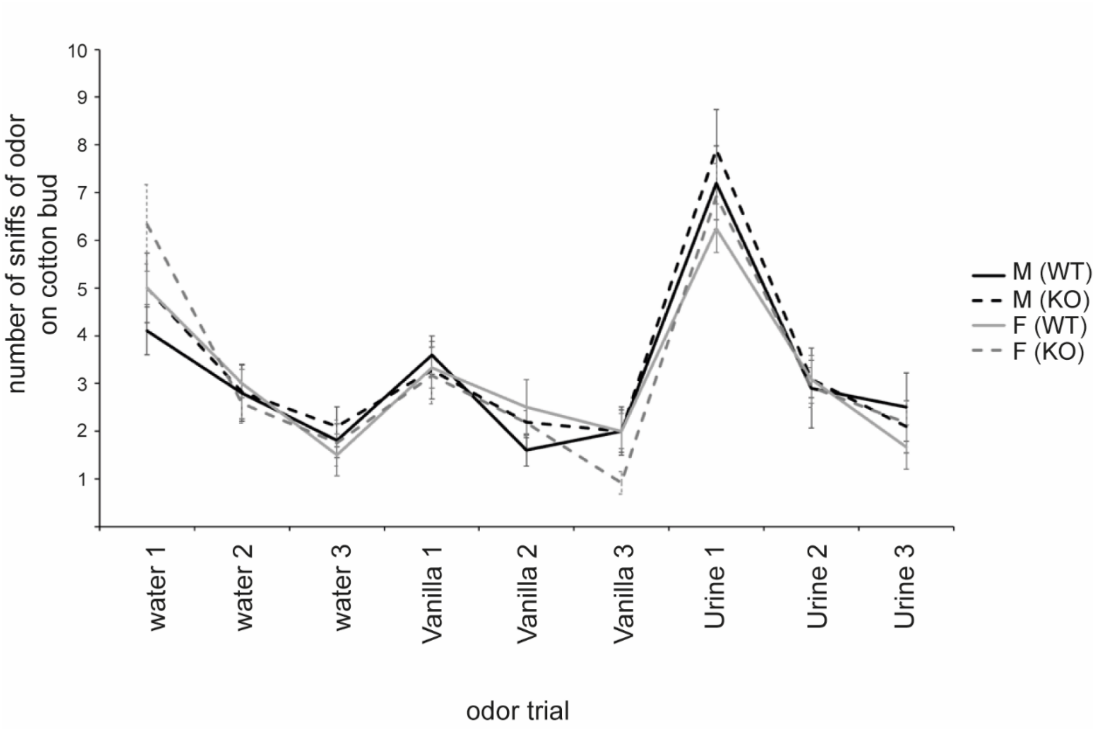


Suppl. Fig 1. Analysis of olfactory habituation/dishabituation in adult *Cntnap2* KO and WT mice of both sexes.

No differences were found in olfactory habituation/dishabituation between KO and WT mice of both sexes, indicating an undisturbed function of the olfactory system. Data presented as means, error bars represent S.E.M. (n=10-12/group).

See Materials and Methods for details.

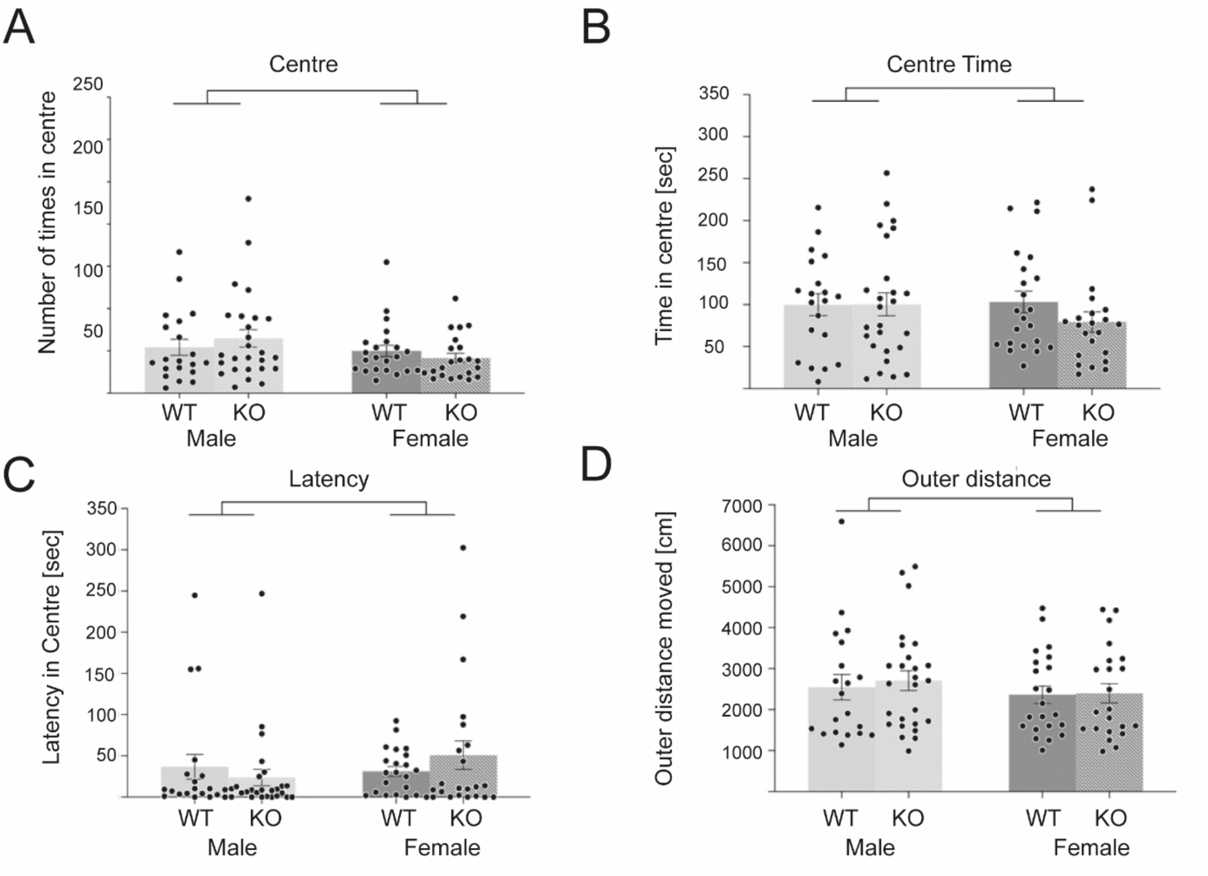


Suppl. Fig. 2. Analysis of anxiety and locomotor activity in male and female juvenile *Cntnap2* KO and WT mice using the Open Field assay.

Avoidance of the centre zone was used as a measure of anxiety (A) and (B), and latency in centre (C) and total distance moved in the outer zone (D) were used as a measure of locomotive activity.

For all parameters, there were no differences between juvenile male WT and *Cntnap2* KO mice. Data presented as means, error bars represent S.E.M. (n>20/group). Statistical analysis was performed using 2-way ANOVA.

See Materials and Methods for details.

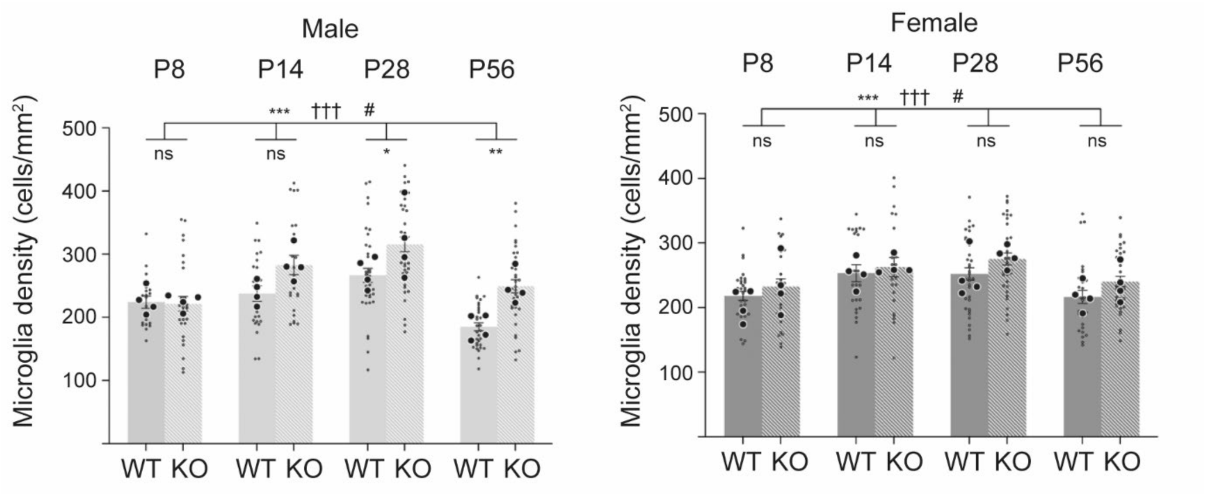


Suppl. Fig. 3. Microglia densities in layer 1 of the ACC of *Cntnap2* KO and WT mice from male and female populations were analysed at four developmental time points P8, P14, P28 and P56.

(A) Microglia cell densities in the male population. There were no differences in cell densities at P8 and P14, but significant increases in KOs compared to WTs at P28 and P56.

(B) In females, there were no significant differences in microglial cell densities at any of the indicated time points.

In all experiments, n=4 KO, n= 4 WT for each timepoint and sex. Large dots represent mice, small dots density on individual sections. Data presented as means, error bars represent S.E.M. Statistical analysis was performed using 2-way ANOVA. Symbols above the bars represent an overall effect of genotype (*), sex (+) or interaction (#), while symbols below the chart represent Tukey's *post-hoc* significance between WT and *Cntnap2* KO (*). In each case 1, 2, or 3 symbols represents $p < 0.05$, $p < 0.01$, or $p < 0.001$.
